# A high-resolution, nanopore-based artificial intelligence assay for DNA replication stress in human cancer cells

**DOI:** 10.1101/2022.09.22.509021

**Authors:** Mathew J.K. Jones, Subash Kumar Rai, Pauline L. Pfuderer, Alexis Bonfim-Melo, Julia K. Pagan, Paul R. Clarke, Sarah E. McClelland, Michael A. Boemo

## Abstract

DNA replication stress is a hallmark of cancer that is exploited by chemotherapies. Current assays for replication stress have low throughput and poor resolution whilst being unable to map the movement of replication forks genome-wide. We present a new method that uses nanopore sequencing and artificial intelligence to map forks and measure their rates of movement and stalling in melanoma and colon cancer cells treated with chemotherapies. Our method can differentiate between fork slowing and fork stalling in cells treated with hydroxyurea, as well as inhibitors of ATR, WEE1, and PARP1. These different therapies yield different characteristic signatures of replication stress. We assess the role of the intra-S-phase checkpoint on fork slowing and stalling and show that replication stress dynamically changes over S-phase. This method requires sequencing on only a single nanopore flow cell, and the cost-effectiveness and high throughput enables functional screens to determine how human cancers respond to replication-targeted therapies.

---

DNA replication stress is characterised by frequent slowing, stalling, and collapse of replication forks which causes unreplicated regions of the genome, DNA damage, mutations, and chromosomal instabilities that drive tumourigenesis (*1, 2*). Many important chemotherapeutic agents target the replication stress response (RSR) pathway (*3*) including inhibitors of ATR (*4*), PARP1 (*5*), WEE1 (*6*), and dNTP synthesis with hydroxyurea (HU) (*7, 8*), as well as DNA crosslinkers like cisplatin (*9*). How these agents affect the movement and fidelity of replication forks across the genome remains unknown due to the low throughput and/or insufficient resolution of current assays. The location of DNA breaks that result from replication fork stalls can be mapped with TrAEL-seq (*10*), but this requires the fork to stall such that it results in a break and this method also carries no information about origin firing or fork movement before the break. DNA combing is a single-molecule method that can measure fork velocity and the frequency of fork stalling (*11, 12*), but the throughput and spatial resolution are both low and fibres cannot be mapped to the genome without resorting to fibre-fish (*13*) which is not scalable to the whole genome. Recently, optical replication mapping (ORM) provided a high-throughput, single-molecule approach to map origin firing genome-wide (*14*), but the 15-kb resolution is too low to detect fork stalling. Long-read sequencing using the Oxford Nanopore Technologies (ONT) platform has enabled the detection of replication origins and fork movement in budding yeast (*15-19*), the malaria parasite *Plasmodium falciparum* (*20*), and human mitochondrial DNA (*21*). Here, we introduce a method that measures fork speed and stalling on single molecules with up to single-nucleotide resolution and use it to show that chemotherapies create different “replication stress signatures”. Our method can clearly distinguish between replication fork slowing and stalling, and by mapping these molecules to the genome, we assess the role of the intra-S-phase checkpoint and show that replication stress dynamically changes over S-phase.

Our DNAscent software can detect two thymidine analogues, BrdU and EdU, on single nanopore-sequenced molecules. When these base analogues are sequentially pulsed into S-phase cells, they are incorporated into the nascent strand by replication forks, leaving a “footprint” of fork movement. Therefore, we pulsed A2058 human melanoma cells expressing PIP-FUCCI (*22*) with EdU, then BrdU, enriched for S-phase cells by fluorescence-activated cell sorting, extracted ultra-high-molecular weight DNA and sequenced it on the Oxford Nanopore MinION platform (Figure 1a-b). As shown in *S. cerevisiae (15, 17, 18*), fork stalling manifests as a sharp drop-off in analogue incorporation into nascent DNA (Figure 1c) and we observed stalled forks in A2058 melanoma cells (Figure 1d). The length (in base pairs) of the base analogue tracks, divided by the 15-minute total pulse length, yielded an accurate measure of fork speed that was consistent between biological replicates (Figure 1e). Our optimised protocol yielded over 180,000 reads with a mapping length greater than 20 kilobases (kb), an N50 of ∼90 kb, and 4,100 called forks from a single MinION flow cell (averages: 150,000 reads longer than 20 kb, 90 kb N50, 2050 fork calls; Supplemental Table S1) with a false positive rate of fork calls less than 0.004% (Supplemental Table S2). This read length and throughput allowed us to capture complex replication dynamics across the entire human genome with multiple forks, origins, and termination sites on the same molecule (Figure 1f).

**Figure 1:**
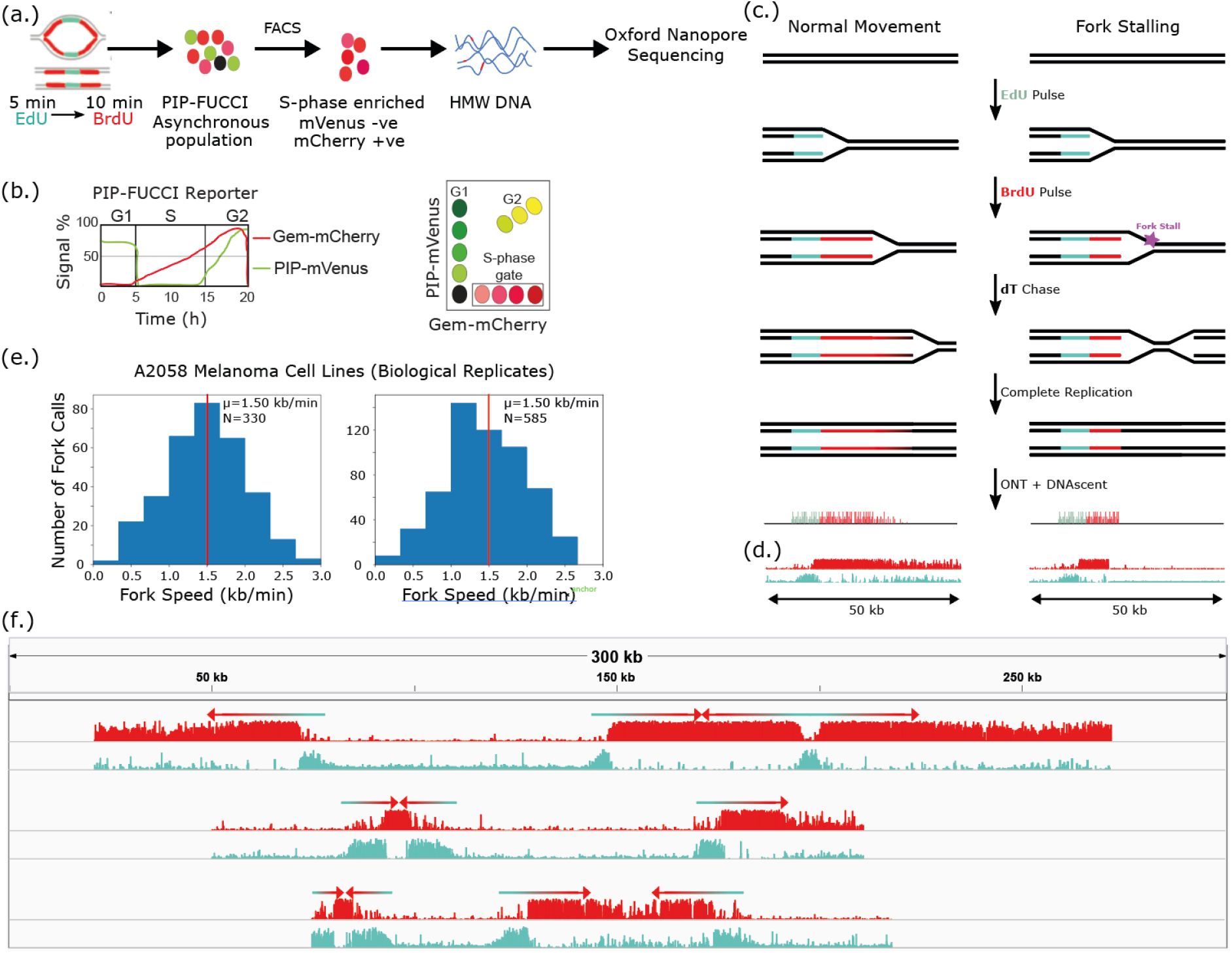
Replication dynamics on ultra-long nanopore reads from human cancer cells. (a.) Cells expressing PIP-FUCCI were sequentially labelled with a 5-minute EdU pulse, a 10-minute BrdU pulse, and a thymidine chase. The asynchronous population of cells was enriched for S-phase cells using FACS (mCherry^+^ mVenus^-^), and high molecular weight (HMW) DNA was extracted and sequenced on the Oxford Nanopore MinION platform. (b.) PIP-FUCCI gates on high levels of geminin (Gem) and low levels of the PCNA-interacting protein (PIP) degron enabling S-phase enrichment without the cellular stress caused by arresting the cell cycle. (c.) Diagram showing that fork stalling manifests as a sudden drop in BrdU incorporation into the nascent strand. (d.) Two nanopore-sequenced molecules from A2058 melanoma cells analysed with DNAscent that show the patterns in panel (c.). Tracks show the probability of BrdU (red) and EdU (blue) at each thymidine position along the molecule. (e.) Distribution of fork speed measured by DNAscent in untreated A2058 cells for two biological replicates. μ, vertical red line: mean; N: number of fork calls. (f.) Three ultra-long single molecules, each represented as a group of two tracks of bar graphs. The top track shows the DNAscent-called probability of BrdU (red) and the bottom track shows the DNAscent-called probability of EdU (blue). Each read is annotated with arrows to show fork direction. Reads were moved onto the same axis from different regions of the genome.

We next treated A2058 cells with either a PARP inhibitor (Olaparib), WEE1 inhibitor (MK1775), ATR inhibitor (VE-821), or HU (Figure 2a). These treatments had a marked effect on replication fork dynamics that was clearly visible on single molecules (Figure 2b). As expected, HU (*12*), ATR inhibition (*23*), and WEE1 inhibition (*24*) all slowed forks (Figure 2c). The dose and treatment time of Olaparib mirrored that of Maya-Mendoza and colleagues (*25*) and, consistent with their findings using DNA combing, we observed a 30% increase in fork speed. Fork speeds from all conditions were consistent across biological replicates (Supplemental Figure S1). To create a quantitative measure of fork stalling, we defined a “stall score” to measure how abruptly BrdU incorporation ends. Stall score is a measure between 0 and 1 of the proportional decrease in the frequency of positive BrdU calls at the BrdU end of the replication fork track. Forks assigned a stall score near 0 are unlikely to be stalled and forks assigned a stall score near 1 are likely to be stalls or pauses (Figure 3a). Stall score provides an additional layer of information about replication stress that does not necessarily track with fork speed: compared to untreated cells, HU and WEE1 inhibition resulted in fork slowing and PARP inhibition resulted in a fork speed increase (see Figure 2c), but all three of these chemotherapies resulted in a similar distribution of stall scores to that of untreated cells (Figure 3b). ATR inhibition did not slow forks as much as HU, but treatment with ATR inhibitors resulted in a marked increase in stall score indicative of frequent fork stalling. While HU is generally thought to rapidly stall replication forks, multiple DNA fibre studies have shown slow but continued fork progression during short and prolonged HU treatment (*26, 27*). Our approach builds on these fibre methods by discriminating between fork slowing in cells treated with HU and rapid stalling in cells treated with ATRi. As before, stall scores from all conditions were consistent across biological replicates (Supplemental Figure S2).

**Figure 2:**
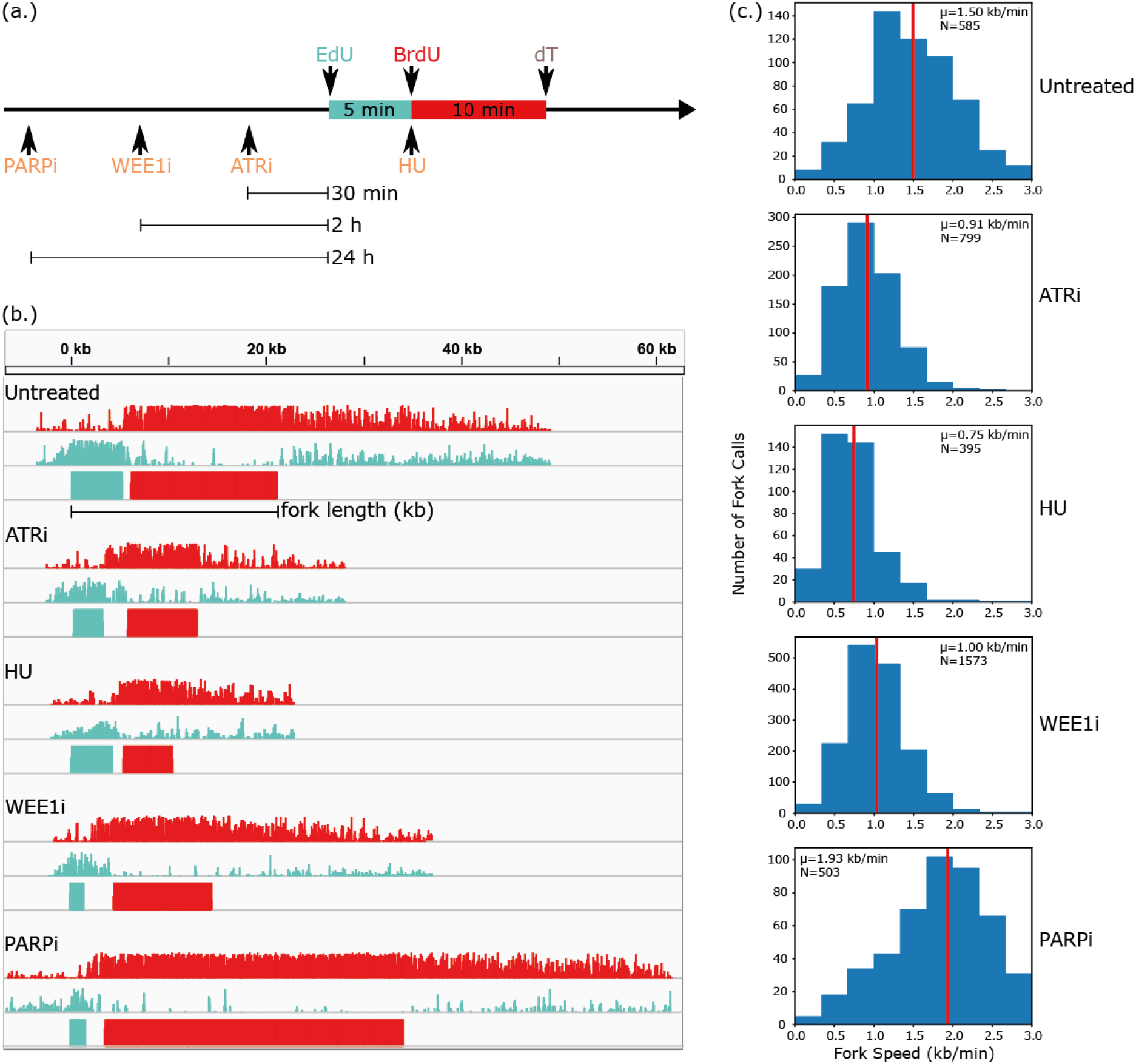
DNAscent measures disruptions to fork speed caused by inhibiting the replication stress response pathway. (a.) A2058 cells were given a monotherapy of either HU, ATR inhibitor, WEE1 inhibitor, or PARP inhibitor at the specified timepoints relative to the BrdU and EdU pulses. (b.) Five single molecules analysed by DNAscent, each showing a rightward-moving fork from untreated cells as well as from each of the monotherapies. Forks were from different regions of the genome, moved onto the same axis so that the start of the EdU tracks align. Each fork is represented as group of three tracks showing (from top to bottom) the probability of BrdU at that position on the read, the probability of EdU at that position on the read, and DNAscent’s segmentation of the read into EdU- and BrdU-positive regions. Fork length was measured by computing the genomic distance, in kilobases, between far ends of the EdU and BrdU segments (shown for the top fork) and dividing this distance by the 15-minute total pulse duration. (d.) Distribution of fork speeds for untreated cells and cells under each monotherapy. Forks near the end of reads, as well as forks together at origins or terminations, were excluded to avoid inaccuracies in fork speed (see Supplemental Methods). μ vertical red line: mean; N: number of fork calls.

**Figure 3:**
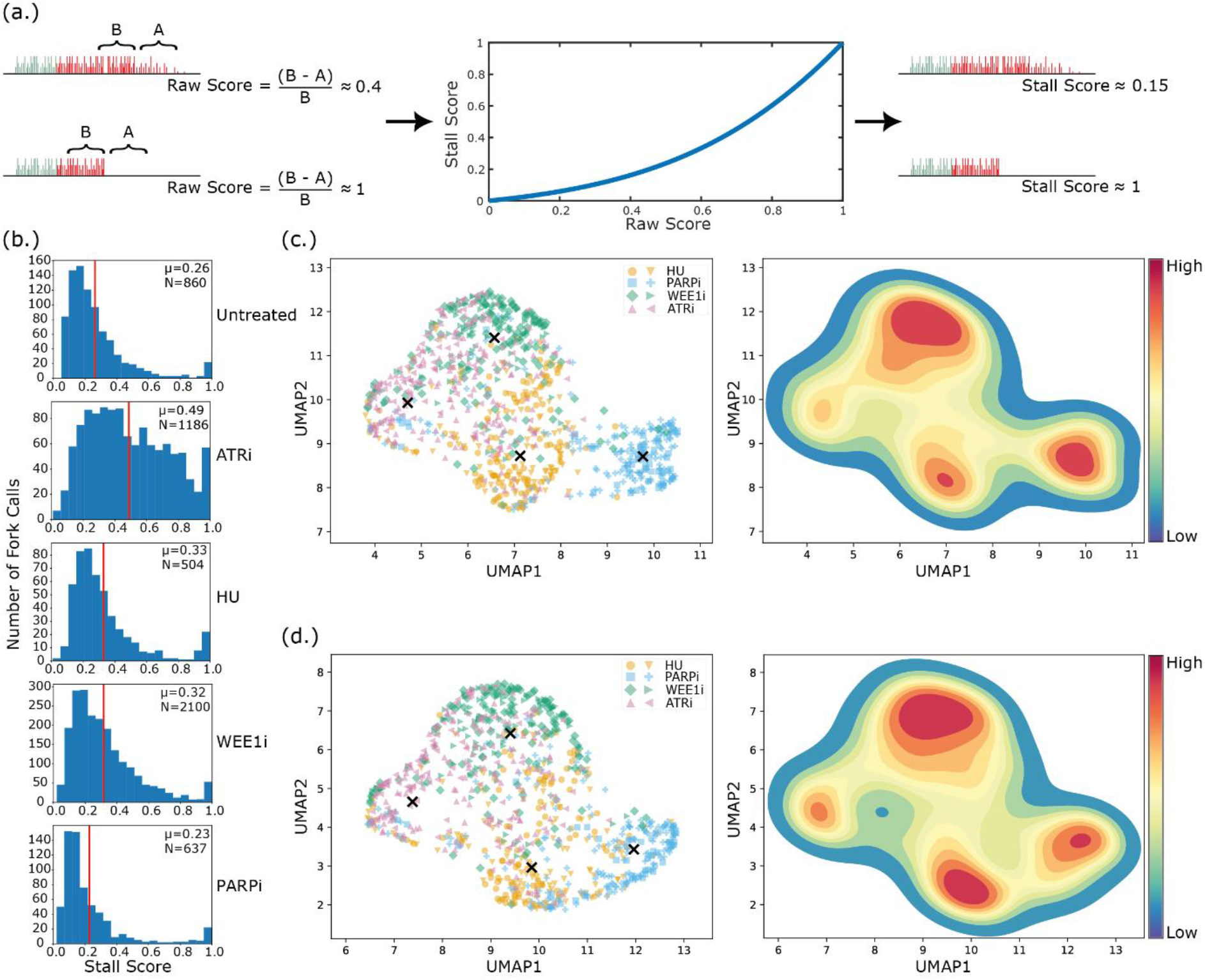
Replication stress signatures stratify cells by treatment. (a.) Each fork is assigned a raw stall score by computing the decrease in frequency of positive BrdU calls (A,B: frequencies) near the end of the fork. A nonlinear scaling is applied to create a final stall score. (b.) Distribution of stall scores for untreated cells, as well as for each monotherapy. μ, vertical red line: mean; N: number of fork calls. (c.) Measurements of the speed, analogue incorporation, and stall score of each fork were used to create an 8-dimensional replication stress signature, shown embedded into two dimensions using UMAP. Left: Each point is the signature for a single fork, points are coloured according to the monotherapy delivered, and biological replicates are shown as different markers. The centroids from K-means clustering (K=4) are shown as black crosses. Right: Kernel density estimation to show distinct clusters, coloured from blue (low density of points) to red (high density of points). (d.) Repeat of panel (c.) that excludes any measurement of fork speed in the replication stress signature.

DNA replication stress is an umbrella term that refers to frequent fork slowing and/or stalling. DNAscent can measure each of these attributes, and to unify them into an overall measure of replication stress, we represented each fork as an 8-dimensional point consisting of features pertaining to fork speed, level of analogue incorporation, and stall score (see Supplemental Methods for details) and reduced points on this 8-dimensional manifold to two dimensions using Uniform Manifold Approximation and Projection (UMAP) (*28*). Forks measured from cells treated with each chemotherapy clustered together, showing that disrupting different elements of the DNA replication stress response (RSR) pathway creates a replication stress signature in this 8-dimensional space (Figure 3c). To demonstrate that our new method captures fundamentally more meaningful information about DNA replication stress than existing methods that just measure fork speed, we repeated the procedure in a 5-dimensional space that excluded any measurement related to fork speed (the track lengths of EdU, BrdU, and the overall fork; see Supplemental Methods). We found that stall score and the level of base analogue incorporation alone were sufficient to create distinct stress signatures of disruption to different elements of the RSR pathway (Figure 3d). These results show that different chemotherapies each create a characteristic replication stress signature.

To investigate whether fork stress changes across S-phase, we applied our method in HCT116 colon cancer cells and HCT116 cells to leverage high-resolution Repli-Seq replication timing data (*29*) and fragile site maps (*30*). We assessed both wild-type HCT116 cells and HCT116 cells with a *CDK2*^AF/AF^ mutation that prevents intra-S-phase checkpoint activation by WEE1 (*31*). While our measured fork speed in wild-type cells was consistent with DNA fibre (*32*), the *CDK2*^AF/AF^ cells showed a slower fork speed and higher stall score than the wild-type (Figure 4a-b). This was consistent with the effect of WEE1 inhibition on fork speed and stress score in A2058 cells, as WEE1 inhibition functionally mirrors a *CDK2*^AF/AF^ mutation (Figure 2c; Figure 3b). We calculated the median replication time in S-phase (Trep) for the genomic position of each called fork using high-resolution Repli-Seq replication timing data for HCT116 cells (*29*) in order to show how the change in fork speed and stall score over S-phase can be measured from a single sequencing run. In wild-type cells, fork speed increased and stall score decreased in genomic positions that tended to replicate later in S-phase (Figure 4c-d). This relationship vanished in the *CDK2*^AF/AF^ mutant, as these cells maintained a constant slow fork speed and high stall score across both early- and late-replicating regions of the genome. In forks mapping to common fragile sites (CFS), we observed a small but significant increase in stall score for wild-type cells, but there was no significant difference between stall score inside and outside of CFS in the *CDK2*^AF/AF^ mutant. While we observed a change in stall score, we did not observe a significant change in fork speed within CFS in either wild-type cells or the *CDK2*^AF/AF^ mutant (Supplemental Figure S3).

**Figure 4:**
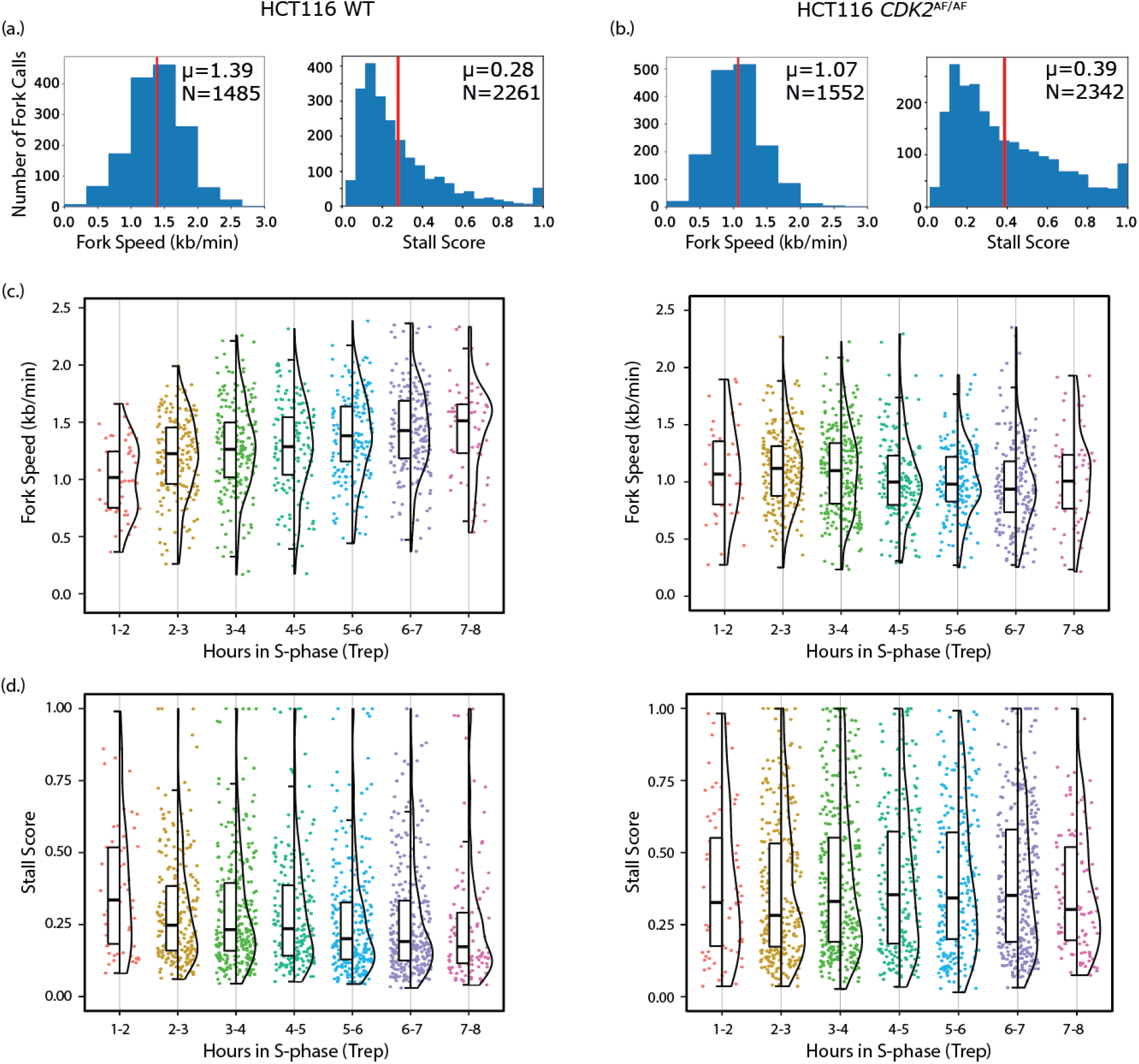
Replication stress dynamics change over S-phase and are checkpoint-dependent. Left column shows results for HCT116 wild-type cells and right column shows results for the HCT116 *CDK2*^AF/AF^ mutant. (a.) Distribution of fork speeds and stall scores for HCT116 wild-type cells. (b.) Distribution of fork speeds and stall scores for the HCT116 *CDK2*^AF/AF^ mutant. For (a.) and (b.), vertical red line shows the mean and the value of the mean (μ) and number of fork calls in the distribution (N) are shown. (c.) Distribution of fork speeds and (d.) stall scores of forks grouped by the median replication time (Trep) of their genomic position. Fork speeds and stall scores mapping to each hour in S-phase are shown as strip plots, violin plots, and box plots with the horizontal black line indicating the median.

We have developed a new method for the high-throughput, high-resolution measurement of DNA replication stress across the human genome; demonstrated that an inexpensive handheld device produces enough data to create replication stress signatures that can stratify cells accurately according to the part of the RSR pathway that was inhibited; and used a single sequencing run to show how replication fork speed changes across replication timing domains. There is significant headroom to improve throughput through future development: The 15-minute analogue pulse represents 3% of an 8-hour human cell S-phase, hence many of molecules sequenced do not include a fork track. A pulldown of EdU prior to sequencing would enrich throughput up to 50x. We have also observed a slight bias towards measuring leading strand synthesis, which may be due to 3’ overhangs on lagging strands reducing capture by transposase during ultra-high molecular weight library preparation. While these are targets for future work, our method significantly supersedes DNA combing in throughput, resolution, cost-effectiveness, automation, and utility and provides additional information about fork dynamics that can be used to quantify fork stress and differentiate the effects of chemotherapies. Replication stress induced by inhibiting different parts of the RSR pathway is not created equally: the impact on replication forks will be highly dependent on the chemotherapy, dose, and timing which will become even more complex when treatments are given in combination. Our method is cost-effective, automated, and provides fundamentally more layers of information to differentiate between stressed forks; all of these attributes are necessary to support high-throughput screens of how cancer subtypes respond to replication-based therapies.

## Methods

### Tissue culture, lentiviral transduction, and S-phase enrichment via FACS

A2058 cells were grown in DMEM with 10% FBS and 1% penicillin–streptomycin. HCT116 cells were grown in McCoy’s 5a modified media with 10% FBS and 1% penicillin–streptomycin. PIP-FUCCI lentiviral transfer plasmid (Addgene: #138714) was cotransfected with psPAX2 pMD2.G into 293T cells. 48 hr later, supernatants were filtered, mixed 1:1 with fresh medium containing polybrene (10 μg/mL), and applied to target cells for 16-24 hr. A2058 and HCT116 cells expressing PIP-FUCCI were labelled with 50 μM EdU for 5 mins, washed 3 x in PBS and labelled with 50 μM BrdU for 10 mins, washed 3x PBS and chased with 100 μM thymidine for 20 mins then trypsinized in 0.05% trypsin. S-phase cells were isolated by FACS sorting (mCherry^+^, mVenus^−^) using a BD FACS Aria Fusion or MoFlo Astrios EQ. Sorted cells were pelleted and stored at −80c. Drug treatments: HU (2 mM), PARPi Olaparib (10 μM), Wee1i MK1772 (1 μM) and ATRi VE-821 (10 μM).

### DNA preparation

UHMW was extracted using Nanobind CBB Big DNA Kit (SKU NB-900-001-01, Circulomics) and UHMW DNA Aux Kit (NB-900-101-01, Circulomics) according to the manufacturer’s protocol (Document ID: EXT-CLU-001). DNA was incubated at 37 °C for 1 h at the elution step. The DNA was eluted in 760 μL Buffer EB (6 million cells for 6x MinION library sequencing) or in 506 μL (4-5.5 million cells for 4x MinION library sequencing). To measure the recovery and the quality of viscus UHMW DNA in Qubit and NanoDrop, the DNA sample was prepared according to the method described by Koetsier and Cantor (*33*). DNA purity was checked by the ratio of DNA concentration measured in NanoDrop and Qubit (1-1.5).

### Library preparation and nanopore sequencing

libraries were prepared using Ultra-Long DNA Sequencing Kit (SQK-ULK001, ONT) and Nanobind UL Library Prep Kit (NB-900-601-01, Circulomics) according to the NanoBind Library Prep - Ultra Long Sequencing protocol (Document ID: LBP-ULN-001; 03/24/2021, Circulomics). UHMW DNA library was eluted in 225 μL (6 million cells) or 150 μL (4-5.5 million cells) ONT Elution Buffer (EB; SQK-ULK001, ONT). The loading library was prepared according to the manufacturer’s protocol and was loaded onto MinION Flow Cell R9.4.1(FLO-MIN106D, ONT). Sequencing run was performed using MinKNOW (v 21.11.7 – 22.08.4) for 96h run script with the sequencing kit set to SQK-ULK001. To load the fresh loading library, the sequencing run was paused after 24 h and the flow cell was washed with Flow Cell Wash Kit (EXP-WSH004, ONT) according to the manufacturer’s guidelines.

### Basecalling and Genome Alignment

Oxford Nanopore sequencing reads were basecalled using Guppy (v5.0.11) using the dna_r9.4.1_450bps_fast configuration. Basecalled reads were aligned to the human genome using minimap2 (v2.17-r941) using the map-ont setting. Reads from A2058 cells were aligned to the chm13v2.0 human reference genome from the Telomere-to-Telomere Consortium (*34*). Reads from HCT116 were aligned to the hg38 (GRCh38.p13) assembly for consistency with the high-resolution Repli-Seq data from Zhao and colleagues (*29*).

### Fork calling with DNAscent

Oxford Nanopore reads were analysed with the new release of our DNAscent software (v3.1.2) available under GPL-3.0 at https://github.com/MBoemo/DNAscent. Each basecalled sequencing run was indexed with the DNAscent index subprogram, and the probability of BrdU and EdU at each nucleotide along each sequenced read was called using the DNAscent detect subprogram. DNAscent detect was run using a minimum mapping quality of 20 and a minimum mapping length of 20 kb. The DNAscent forkSense subprogram segmented each read into EdU- and BrdU-positive regions. These were matched into replication forks, such that the replication fork track ends where BrdU incorporation starts to decrease at the start of the thymidine chase. Fork calls on each read were then matched into origins and termination sites.

### Measurement of fork speed with DNAscent

DNAscent forkSense outputs replication fork calls as bed files indicating the region of the genome each called fork moved through during the 15-minute EdU-BrdU pulse. The length of each region (in kb) was divided by 15 minutes to compute the speed of each fork in kb per minute. These fork tracks will be shorter if (i.) the forks are from an origin that fired during the pulse, (ii.) the forks come together in a termination site during the pulse, or (iii.) the fork track runs off the end of the sequenced molecule. To present an accurate picture of fork speed, fork tracks were excluded from the analysis if they were on a read with a called origin or termination site, or if they started or ended within 3 kb of the end of the read.

### Fork stall calling with DNAscent

The number of positive BrdU calls (probability > 0.5 called by DNAscent detect) was calculated over a 2 kb window before (inside the BrdU-positive segment) the BrdU end of fork track and divided by the number of thymidine bases in the 2 kb window resulting in a fraction between 0 and 1 (denoted *B*). This fraction of positive BrdU calls divided by attempts was also calculated over a 2 kb window immediately after (outside the BrdU-positive segment) the BrdU end of the fork (denoted *A*). A raw score was computed:

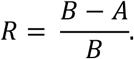

A nonlinear scaling was applied to the raw score *R* to compute a stall score between 0 and 1:

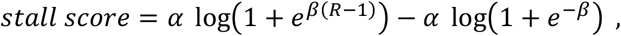

where α=1.55 and β=3. This scaling is plotted in Figure 3a and serves a dual role. First, it accounts for the fact that a fork which has not stalled during the analogue pulse will produce a relatively high raw score approximately between 0.2 and 0.4 and rescales this range closer to zero. Second, it creates more conservative estimates of raw scores in the 0.6-0.9 range so that only high-confidence fork pauses or stalls have stall scores near 1. Each fork call is annotated with a stall score in DNAscent v3.1.2. To avoid erroneous results, DNAscent declines to assign a stall score to forks where (i.) the fork track runs off the read, (ii.) forks that come together in a termination site during the pulse, and (iii.) forks where there is an insertion or deletion longer than 100 bp in the read inside one of the 2 kb windows. These forks were excluded from the analysis in Figures 3-4.

### Replication stress signatures with DNAscent

The following eight features were measured for each fork: (i.) the total length of the fork track (in bp), (ii.) the length of the EdU track (in bp), (iii.) the length of the BrdU track (in bp), (iv.) fraction of thymidine positions called as BrdU in the EdU segment, (v.) fraction of thymidine positions called as EdU in the EdU segment, (vi.) fraction of thymidine positions called as EdU in the BrdU segment, (vii.) fraction of thymidine positions called as BrdU in the BrdU segment, (viii.) the stall score. DNAscent v3.1.2 outputs these features as a bed file for ease of use. Each of these eight features, across all forks, was rescaled to the interquartile range and reduced to two dimensions using Uniform Manifold Approximation and Projection (UMAP)(*28*) with 250 nearest neighbours, a minimum embedded point distance of 0, and the Chebyshev metric. A total of 125 forks from each sequencing run (a total 250 forks per monotherapy, accounting for two biological repeats) were used. Kernel density estimates were made using Seaborn (v0.11.2) with the default settings.

### Mapping forks to Trep and CFS

Gaussian smoothed and scaled high-resolution Repli-Seq data for HCT116 cells was accessed from NCBI GEO (accession number GSE137764) and Trep was calculated via sigmoid fitting as per the authors’ instructions(*29*) for each 50 kb bin across the human genome. The Trep for each DNAscent fork call was computed using the Trep of the 50 kb genomic region that the fork call mapped to. Common fragile site (CFS) locations were accessed from the database from Kumar and colleagues (*30*).

## Acknowledgements

The authors would like to thank Francis Totañes and Catherine Merrick (Department of Pathology, University of Cambridge), Brian Gabrielli (Mater Research Institute), Valentine Murigneux (UQ-GIH), and all the members of the Jones and Clarke laboratories (Diamantina Institute) as well as the Boemo laboratory (Cambridge Pathology) for guidance and helpful conversations. We would like to acknowledge the contribution of David Sester, Dalia Khalil, Yitian Ding and Andy Wu at the Translational Research Institute (TRI) Flow Cytometry Facility for invaluable help and technical assistance. This research was carried out in part at the Translational Research Institute, Australia. The Translational Research Institute is supported by a grant from the Australian and Queensland Governments

## Author Contributions

MJKJ, SEM, and MAB conceptualised and designed the study. MAB and PLP developed the software and did the bioinformatics analyses. MJKJ and SKR conducted the wet lab research and collected the data. JKP provided reagents and intellectual contributions. MJKJ, MAB, and PRC acquired funding for the project. MAB, MJKJ, and A.B-M. made the figures. MJKJ and MAB supervised the project and wrote the manuscript with input from all authors.

