## Supplemental Information for "A high-resolution, nanopore-based artificial intelligence assay for DNA replication stress in human cancer cells"

### Supplemental Figures

**Figure S1: Measured replication fork speed is consistent across biological replicates.** The distribution of fork speeds are shown for two biological replicates (columns) of each treatment (rows). The left column is shown in Figure 2c of the main text. Vertical red line shows the mean, and the value of the mean (μ) and number of fork calls in the distribution (N) are shown for each replicate.

**Figure S2: Measured stall score is consistent across biological replicates.** Similar to supplemental Figure S1, but showing the distribution of stall scores for two biological replicates (columns) of each treatment (rows).

**Figure S3: Fork speed and stress measurements in common fragile sites (CFS) for HCT116 cells.** Distribution of replication fork speeds (top row) and stall scores (bottom row) for replication forks that mapped inside and outside common fragile sites (CFS) for HCT116 wild-type cells (left) and HCT116 *CDK2*$^{AF/AF}$ (right). Quartiles are shown as inner box plots (dark grey) and medians are shown as white dots. The difference in stall score for forks that mapped inside versus outside CFS was significant for wild-type cells (Mann-Whitney U test, $p<0.05$).

**Figure S4: Sorting for S-phase cells using PIP-FUCCI.** Representative plot of PIP-FUCCI sorting (A2058 cell lines treated with Olaparib for 24 hours) showing the S-phase gate (red box).

### Supplemental Tables

**Table S1: Yields from Oxford Nanopore and DNAscent from each sequencing run**. The number of reads shown for each sequencing run is the number of reads that had a mapping length to the reference genome greater than 20 kb, a mapping quality greater than or equal to 20, and passed the quality controls in DNAscent v3.1.2. The N50, mean read length, and median read length were all calculated on this group of reads rather than on all reads from the sequencing run, as short reads will not be long enough to measure replication fork speed and stress given the 15-minute BrdU-EdU pulse used.

**Table S2: False positive rates for human genomic DNA not treated with BrdU or EdU**. Each row is a biological replicate, and the number of reads shown for each sequencing run is the number of reads that had a mapping length to the reference genome greater than 20 kb, a mapping quality greater than or equal to 20, and passed the quality controls in DNAscent v3.1.2.

Supplemental Figure S1

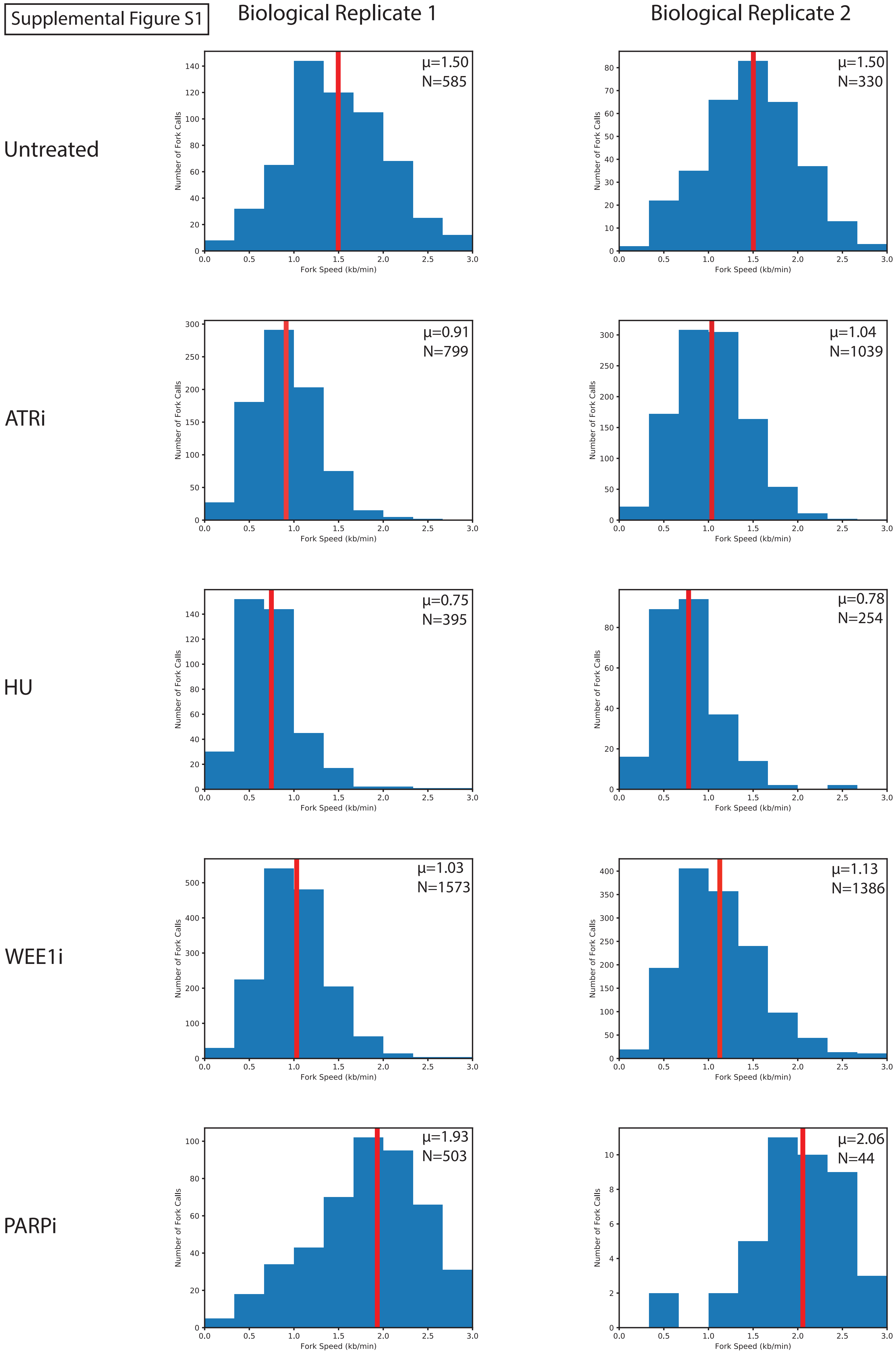

Supplemental Figure S2

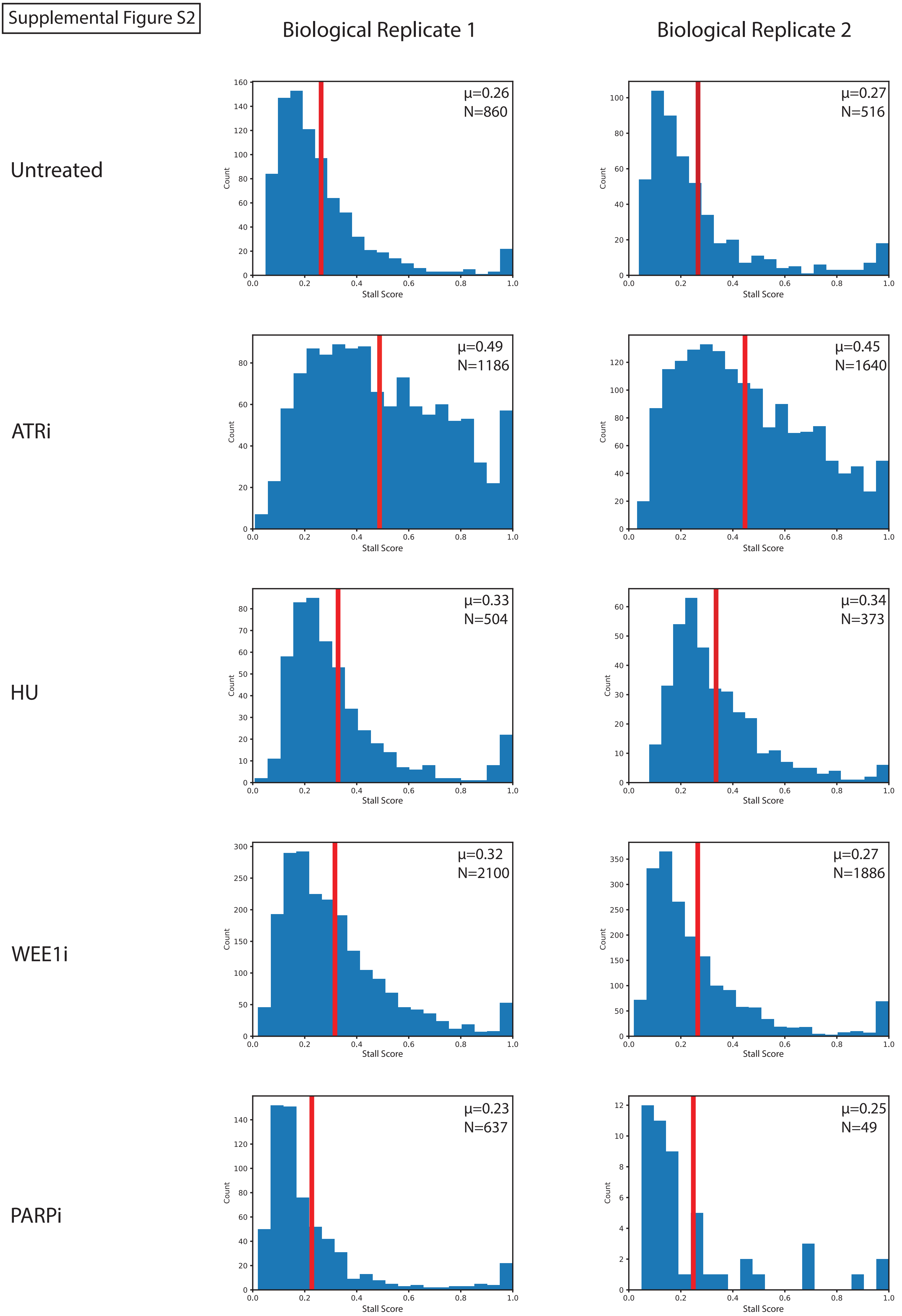

Supplemental Figure S3

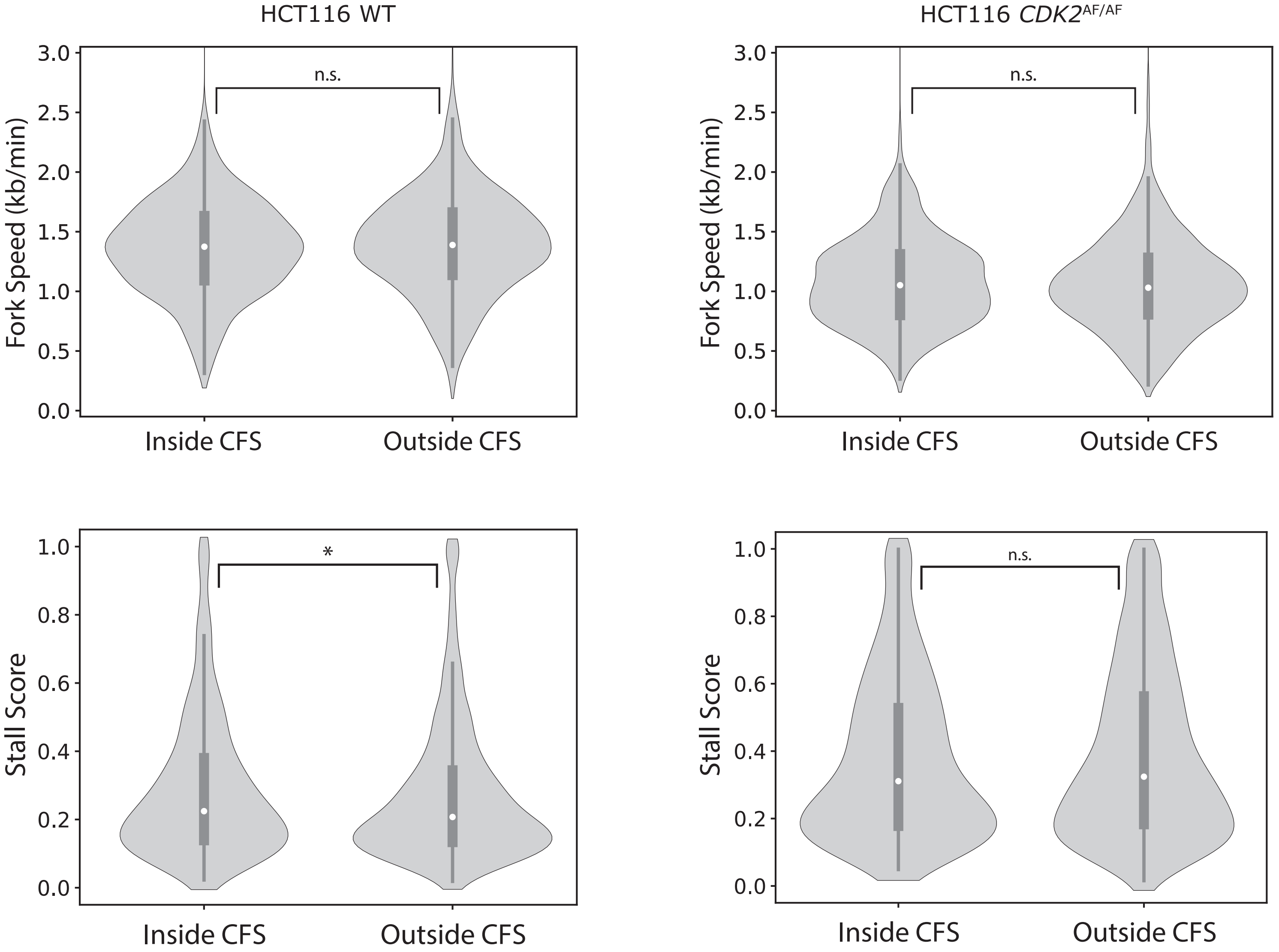

Supplemental Figure S4

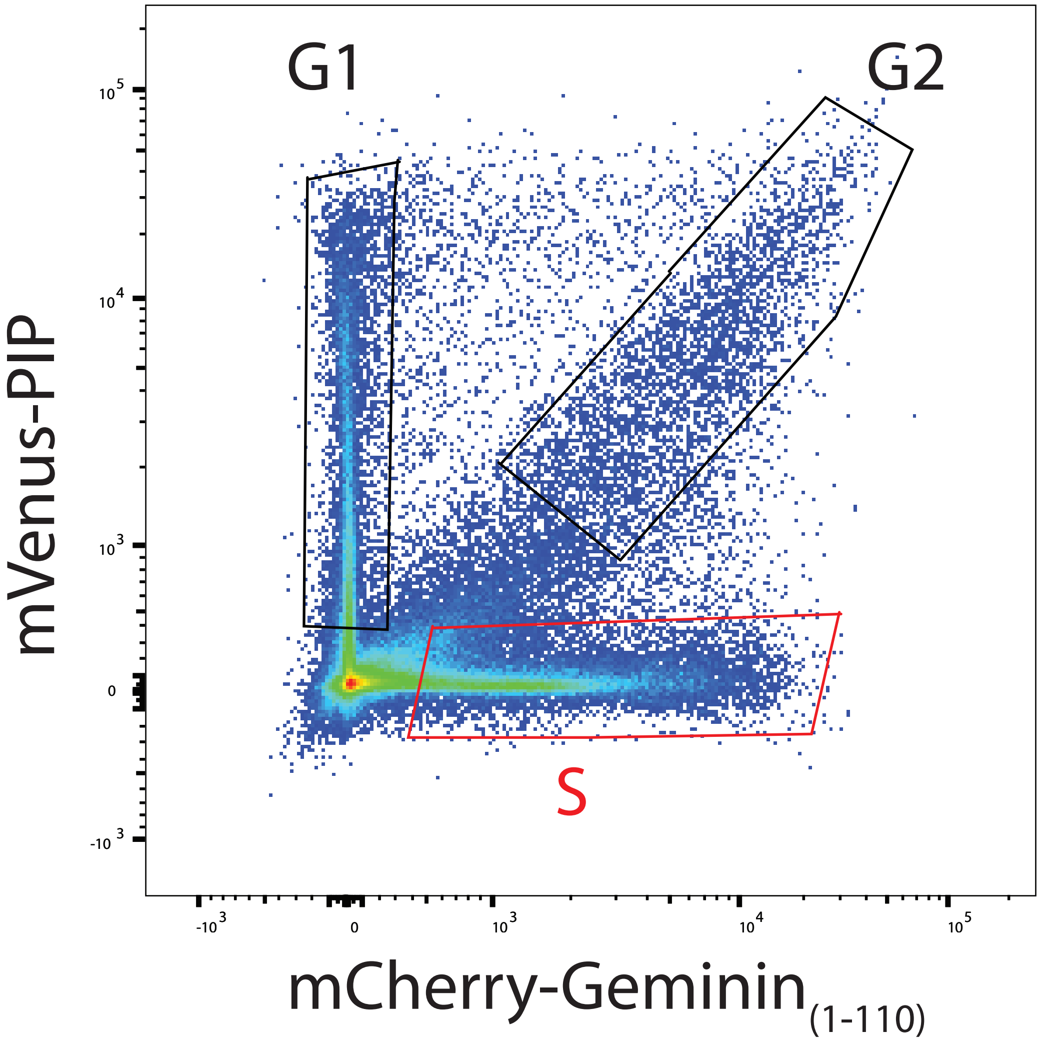

Supplemental Table S1

| Cell Type/Treatment | Number of reads | N50 (kb) | Mean Read Length (kb) | Median Read Length (kb) | Left fork calls | Right fork calls | Origin calls | Termination calls |
|---|---|---|---|---|---|---|---|---|
| **A2058-Untreated** | 60900 | 88.5 | 67.8 | 50.3 | 484 | 511 | 103 | 76 |
| **A2058-Untreated** | 133553 | 84.7 | 66.2 | 50.5 | 848 | 845 | 106 | 101 |
| **A2058-ATRi** | 146965 | 85.8 | 67.0 | 51.4 | 925 | 906 | 180 | 134 |
| **A2058-ATRi** | 216198 | 91.5 | 70.3 | 53.0 | 1275 | 1264 | 279 | 185 |
| **A2058-HU** | 35485 | 91.2 | 70.0 | 52.8 | 241 | 259 | 41 | 47 |
| **A2058-HU** | 108923 | 99.4 | 74.2 | 54.6 | 501 | 514 | 44 | 57 |
| **A2058-WEE1i** | 175683 | 98.0 | 73.3 | 54.0 | 1597 | 1672 | 228 | 213 |
| **A2058-WEE1i** | 228003 | 82.9 | 65.4 | 50.8 | 1774 | 1760 | 210 | 187 |
| **A2058-PARPi** | 60063 | 92.9 | 70.5 | 52.2 | 40 | 54 | 2 | 4 |
| **A2058-PARPi** | 262244 | 98.5 | 74.2 | 55.1 | 607 | 611 | 42 | 39 |
| **HCT116-WT** | 181241 | 89.2 | 68.7 | 51.5 | 2051 | 2135 | 379 | 363 |
| **HCT116-*CDK2*$^{AF/AF}$** | 199360 | 86.7 | 66.9 | 49.5 | 2018 | 2016 | 452 | 350 |

Supplemental Table S2

| Cell Type/Treatment | Number of reads | EdU region calls | BrdU region calls | Left fork calls | Right fork calls | Origin calls | Termination calls |
|---|---|---|---|---|---|---|---|
| **RPE1-No BrdU or EdU** | 4151 | 0 | 0 | 0 | 0 | 0 | 0 |
| **RPE1-No BrdU or EdU** | 106716 | 3 | 4 | 2 | 2 | 1 | 0 |
